# Characterization of Cell-Induced Astigmatism in High-Resolution Imaging

**DOI:** 10.1101/2021.10.01.462719

**Authors:** Rick Rodrigues de Mercado, Hedde van Hoorn, Martin de Valois, Claude Backendorf, Julia Eckert, Thomas Schmidt

## Abstract

High-resolution and super-resolution techniques become more frequently used in thick, inhomogeneous samples. In particular for imaging life cells and tissue in which one wishes to observe a biological process at minimal interference and in the natural environment, sample inhomogeneities are unavoidable. Yet sample-inhomogeneities are paralleled by refractive index variations, for example between the cell organelles and the surrounding medium, that will result in the refraction of light, and therefore lead to sample-induced astigmatism. Astigmatism in turn will result in positional inaccuracies of observations that are at the heart of all super-resolution techniques. Here we introduce a simple model and define a figure-of-merit that allows one to quickly assess the importance of astigmatism for a given experimental setting. We found that astigmatism caused by the cell’s nucleus can easily lead to aberrations up to hundreds of nanometers, well beyond the accuracy of all super-resolution techniques. The astigmatism generated by small objects, like bacteria or vesicles, appear to be small enough to be of any significance in typical super-resolution experimentation.

## 1. Introduction

High-resolution and super-resolution optical microscopy is routinely used in biology, biophysics and material science. For homogeneous samples a lateral resolution down to 10 nm and an axial resolution down to 20 nm has been demonstrated [1]. At this resolution it was possible to discover among others the cytoskeletal organization that structures the neuronal axon [2] and the dynamics of clathrin-mediated endocytosis in live cells [3]. In most of the discoveries the studied objects were either very thin (< 500 nm) and positioned close to the cover slip, or the embedding medium was largely homogeneous. The optical resolution in thicker samples, or even tissue significantly decreases [4]. On one side the resolution decrease is due to an increase of scattering and background signal. On the other side in thick, optically inhomogeneous samples, inhomogeneities induce local astigmatism that leads to local image blur. Such blur might not be visible for regular, diffraction-limited imaging, yet it affects images acquired in super-resolution modality [5–7].

In our studies we are concerned with traction force measurements to study cell mechanics [8–11]. We use flexible micropillar arrays [12] on which cells are grown and imaged. Cells and arrays are imaged in the upside-down configuration on an inverted microscope (see Fig.1A, left) that allowed us to determine the cellular traction forces and simultaneously image sub-cellular structures at super-resolution modality [13]. Deflections of pillars from their equilibrium position thereby report on the forces that cells apply.

**Fig. 1.**
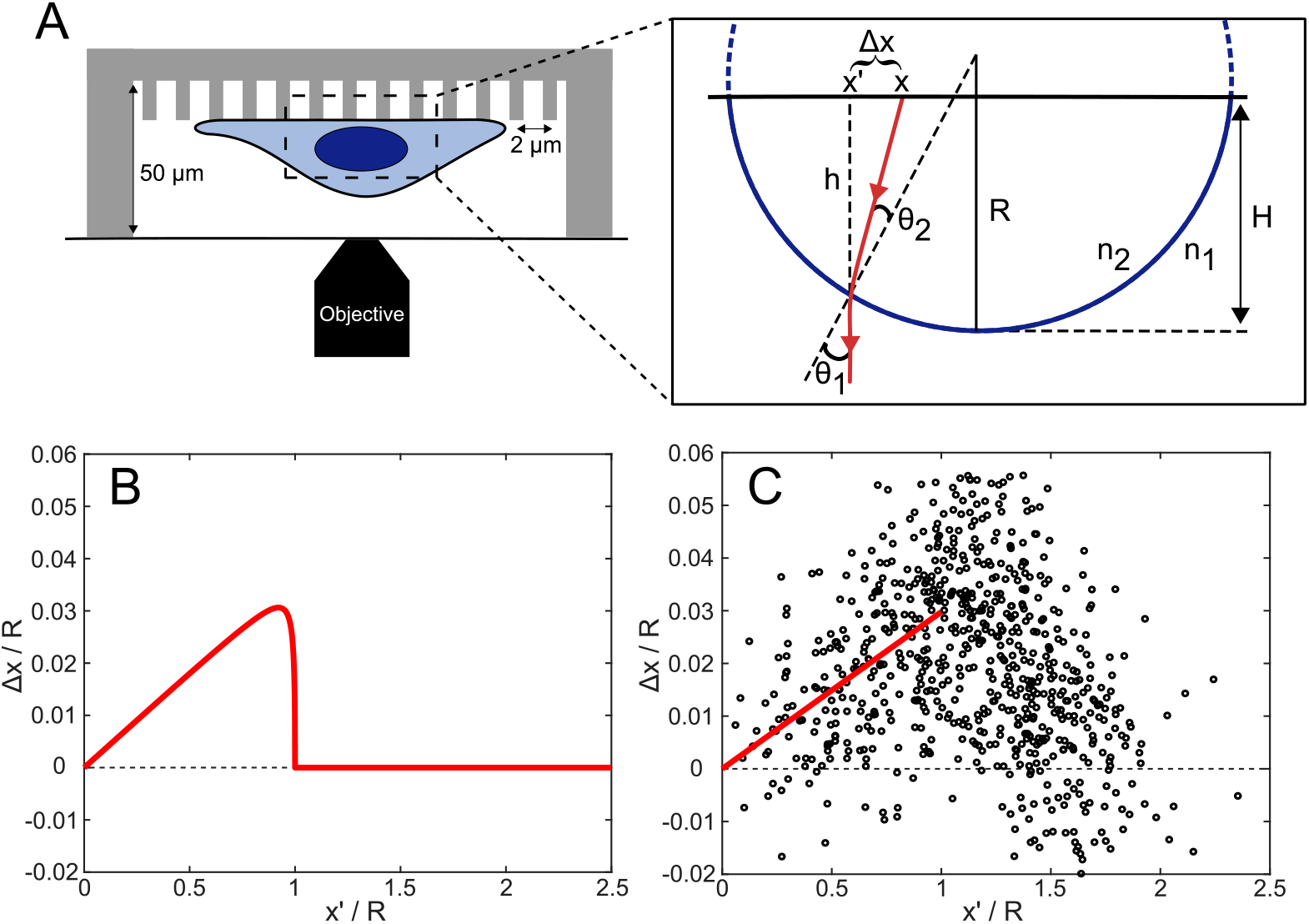
Refraction at the nucleus-cytosol interface causes apparent deformations beyond the accuracy of super-resolution microscopy. **A.** Schematic drawing of a micropillar array mounted onto an inverted microscope (left) and a theoretical schematic of a light-ray refracting at the nucleus-cytosol interface resulting in an apparent deflection Δ*x* (right). **B.** Theoretical prediction of refraction at a spherical surface for values typically relevant for cells (*R* = 10 μm, *H* = 7 μm, *n*_1_ = 1.33 and *n*_2_ = 1.38). **C.** Displacement of dots underneath 16 metaphase cells on a 2.5 MPa flat layer of PDMS. A fit of Eq.(2) for *x′* < *R* is shown in red, where *H*/*R* · (*n*_2_/*n*_1_ – 1) = 0.0298 ± 0.002.

Our modality faithfully allowed us to image structures and measure forces at the thin perimeter of cells. Yet, in thicker samples imaging artifacts limit the resolution. The refractive index of cells differs from the surrounding medium and even within cells, the index is not constant [14]. This holds in particular for larger cellular objects like the nucleus. The refractive index change induces refraction of light at the cell-medium or the nucleus-cytosol interface. This refraction poses a serious problem when measuring traction forces using micropillar arrays. The position of the pillars is measured through the cell. Refraction leads to an erroneous position estimation that is at the heart of most super-resolution techniques.

Here we determine the cell-induced astigmatism during traction force measurements on micropillar arrays. We introduce a simple model that explains the observed distortions of the substrate as a lensing effect of the cell nucleus. The model permits to quickly assess the impact of astigmatism as a useful estimator for the predicted image quality in a given sample. The estimator may be applied to any optical measurement technique.

We tested the model for HeLa cells undergoing mitosis. The astigmatism changed as mitosis progressed and was dependent on the location and the shape of the cell nucleus. As predicted, astigmatism vanished at the center of the nucleus and increased to up to 400 nm at its edge. The astigmatism was observable well beyond the outline of the nucleus. Based on our model we found an estimation for the refractive index of the nucleus of ~ 1.369.

## 2. Materials and methods

### 2.1. Cell culture

HeLa cells were cultured in high-glucose Dulbecco’s Modified Eagle’s Medium (DMEM, Sigma-Aldrich, D1145) supplemented with 10% fetal calf serum (Thermo Scientific), 2 mM glutamine and 100 μg/mL penicillin/streptomycin at 37 °C with 5% CO_2_. We utilized cells transformed with H2B-GFP, a construct that labels the histone 2B protein with a green-fluorescent protein. Prior to measurements, cells were synchronized in late G2 phase using the cell-cycle blocker RO3306 (Sigma-Aldrich, SML0569) [15]. This inhibitor was added to the cell medium in a final concentration of 10 μM. Cells were seeded at single cell density directly on the micropillar array or flat layer of PDMS and allowed to spread ~ 16 - 20 hours. Cells were released from the RO3306 block by refreshing the medium. Substrates were subsequently inverted onto #0, 25 mm diameter, round coverslips (Menzel Glaser). Spacers of 50 μm height prevented cells from touching the coverslip. Measurements were done shortly after the release.

### 2.2. Micropillars and flat PDMS layers

Hexagonal-arranged arrays of poly-di-methyl-siloxane (PDMS, Sylgard 184, Dow Corning) micropillars of 2 μm diameter, 2 μm spacing and with a height of 4.1 and 6.9 μm were produced using replica-molding from a silicon wafer into which the negative of the structure was etched by deep reactive-ion etching. Short and tall micropillars on the array had a characteristic bending modulus of 47.2 and 9.8 kPa and a stiffness of 65.9 and 13.7 nN/μm, respectively. The pillar arrays were flanked by integrated 50 μm high spacers (as described in [16] and schematically shown in Fig. 1A) such that pillar tops and hence cells attaching to them were within the limited working distance of our high-NA objective (<170 μm) on an inverted microscope. The use of a high-NA objective is a prerequisite for any high-resolution optical imaging. The micropillar arrays were kept from floating using a support weight made from glass. The temperature was kept at 37 °C with constant 5% CO_2_ concentration in a stage-top incubator (Tokai Hit, Japan). Flat PDMS layers with a Young’s modulus of 2.5 MPa were produced by molding on the flat parts of the silicon wafer. During measurements, flat layers were placed in upside down position and supported on a 50 μm spacer such that the spacing was similar to the micropillar conditions.

The tops of the micropillars were coated with a mixture of Alexa568-labeled and unlabeled fibronectin (Sigma-Aldrich, F1141) (ratio 1:5) using micro-contact printing. A 40 μL drop with 60 μg/mL fibronectin was incubated 1 hour on a flat piece of soft PDMS (1:30, crosslinker:base ratio), washed with ultrapure water and left to dry under laminar flow. After 10 minutes of UV-Ozone (Jelight) activation of the micropillars, micro-contact printing was performed for 10 minutes. Non-printed areas were blocked 1 hour with 0.2% Pluronic F-127 (Sigma, P2443) in PBS.

Flat PDMS layers were prepared similarly. A printed micropillar array was used to transfer the hexagonal pillar pattern in 10 minutes onto a flat layer of PDMS, which was treated with UV-Ozone for 10 minutes. Subsequent blocking of unlabeled areas was performed as above. The position of the pillar tops was observed by fluorescence microscopy at 561 nm excitation (supplementary Fig. S2).

From those fluorescence images, the exact pillar-centroid positions were determined down to 30 nm accuracy. The deflection precision of 30 nm corresponded to a force accuracy of 2 and 0.4 nN for the short and tall pillars, respectively. All analysis was done using specifically designed software (MATLAB, Mathworks).

### 2.3. Microscopy

Imaging was performed on an inverted microscope (Zeiss Axiovert 200) with a 100X, 1.4 NA oil objective (Zeiss). The setup was expanded with a Confocal Spinning Disk unit (Yokogawa CSU-X1), an emCCD camera (Andor iXon DU897) and a home-built focus-hold system at the rear port. The micropillar arrays were imaged on 561 nm laser illumination (Cobolt). A 488 nm laser (Coherent) was used to image the chromosomes (H2B-GFP). Images of the chromosomes were captured in a single plane 5 μm below the plane defined by the tops of the micropillars.

The cell-cycle phase was quantified comparing the shape of the nucleus to the predicted visual state given by literature [17, 18].

### 2.4. Analysis

Image analysis was performed using specifically designed MATLAB scripts. Pillar-top localization was performed as described previously [16]. A cell-mask was created by dilation of 10 μm of a threshold mask of the nucleus. Deflections within this mask were decomposed in radial and tangential components relative to the center of the nucleus. The radial component of the deflection was much larger as compared to the tangential deflection (Fig. S3). Relative to the radial deflection center(s), inward deflections were defined as negative radial deflections, while outward deflections were defined as positive deflections (see supplementary Fig. S4). In each image, the absolute deflection of pillars within the cell mask were compared to the background. Only deflections beyond the background were used when assessing the mean deflection per pillar. Dunn’s test pairwise comparison after a Kruskal-Wallis test was used to determine statistical significance between populations. Data sets were significantly different with probabilities of *p* < 0.001 (***) and not significantly different with probabilities of *p* > 0.05 (ns).

## 3. Results

### 3.1. Theoretical model: nuclear lensing induces large optical aberrations

At the interface between two media of different refractive indices, refraction occurs. The curvature of the interface and the refractive index difference determine the strength of the effect which is, since Huygens, used to make lenses [19]. Any curved object, including cells, will display lensing. In particular, the cell nucleus as having a different refractive index [14] when compared to the cytosol and to the culture medium is the major source of cellular lensing (Fig.1 A). Since the cell shape changes considerably from flat to round during the cell cycle, the lensing effect is dynamic on the about 2 hour timescale for cell division. Here, we present a simple model that explains the apparent deformations of the substrate below the cell due to the refraction of light at the interface between the nucleus and its surrounding.

The nucleus of refractive index *n*_2_ was modeled as a spherical cap of radius *R* and height *H* embedded in a medium of refractive index *n*_1_ (Fig. 1 A, right). Rays passing the object are refracted leading to an imaginary lateral displacement Δ*x* that depends on the distance of the object from the center of the cap *x*. In the model we assumed that rays are normal to the image plane, a realistic assumption for the case of microscopy, where the focal distance of the imaging lens (≈ 10 mm) is much larger than the radius of the nucleus (≈ 10 μm). In radial direction the displacement is given by (see also the supplementary information)

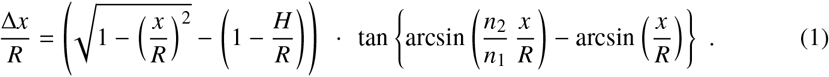

For typical values relevant to cells (*R* = 10 μm, *H* = 7 μm, *n*_1_ = 1.33 and *n*_2_ = 1.38) aberrations close to the edge of the spherical cap can be as large as ≈ 400 nm, close to or beyond the diffraction limit in regular microscopy, yet significant larger than the typical resolution in super-resolution microscopy (Fig. 1B). For positions close to the center, i.e. *x* ≪ *R*, eq. (1) reduces to

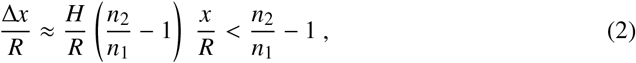

displaying a linear increase of the lateral displacement with distance from the center. For the typical values of the refractive indices mentioned above, the relative displacement is smaller than Δ*x*/*R* < 0.038.

Given the situation sketched in Fig.1 we here define an easy estimator that allows to assess whether astigmatism is relevant to a given imaging mode. Seeking a resolution of *δx*, the imaginary displacement, as obtained from Eq.(1) or Eq.(2), should not exceed the value of

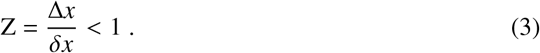

In regular microscopy, resolution is determined by the diffraction limit, *δx*_reg_ = *λ*/(2 NA). For a typical wavelength of *λ* = 600 nm, and numerical aperture, NA= 1.4, the *Z* < 1 for an object (nucleus) with a radius of *R* < 5.7μm. For super-resolution microscopy the achieved accuracy typically is *δx* = 30 nm, which hence limits the typical object size to about *R* < 0.8μm, much smaller than the typical size of the nucleus of a mammalian cell.

### 3.2. Experimental verification: nuclei cause significant distortions of nearby substrates

Experiments were performed to verify the above predictions, namely the apparent deformation of the substrate below cells caused by the lensing effect of the nuclei. We found that refraction of light leads to distortions that follow the behavior in our model, reaching values of up to 400 nm. Cells were grown on a flat substrate that was prior stamped with a micrometer-spaced highly regular hexagonal pattern of labeled fibronectin. The sample was mounted in the upside-down configuration onto the microscope such that the pattern was imaged through the cells. Subsequently, the center-of-mass position of all dots of the pattern was analyzed and compared to the nominal spacing set by the nano-fabricated master previously verified by EM imaging [16].

The radial deflections of 725 dots below or next to 16 metaphase cells on the stiff, flat layer were combined in Fig.1 C. The relative displacements to the nominal positions Δ*x*/*R*, were plotted against the relative distance from the center of the nucleus *x′*/*R*. Two regimes were distinguished, based on the distance from the center of the nucleus and the size of the nuclei *R*. At the center of the nucleus, the deflection was close to zero. For dots located below the nuclei (*x′* < *R*) the deflections linearly increased with distance from the center. For dots surrounding the nuclei (*x′* > *R*) the deflections decreased steeply with increasing distance from the center of the nucleus, returning to zero near the cell edge. The maximal relative deflection we determined, which was localized close to the edge of the nucleus, increased up to Δ*x*/*R* = 0.05.

The experimental observations follow our model predictions. The radial deflection increased linearly as pillars were closer to the edge of the nucleus. Deflections decreased rapidly outside the nucleus. It should be noted that most of the deflections were beyond the localization accuracy of 30 nm that is typical for high-resolution imaging, and hence *Z* > 1. Eq.(2) was subsequently used to determine the refractive index of the cell’s nucleus. As cells rounded up significantly during metaphase, we used that *H*/*R* ≈ 1 (Fig. 3). Using this approximation we found a value of *n*_2_ ≈ 1.369. This result corroborates values obtained by other methods earlier [14].

### 3.3. Effect of astigmatism on traction force measurements

In a followup experiment we resorted to experiments in which interphase cells were grown on soft (9.8 kPa) micropillar arrays. It is known that the height of interphase cells is typically low, cells spread out significantly, their peripheral cytosol is very thin and that interphase cells mainly pull on the substrate from their periphery [20, 21]. Fig. 2A shows a HeLa cell that was attached to a pillar array pulling on the fibronectin-coated pillar tops. Pulling of the cell resulted in the displacement of individual pillars in the image (white arrows in Fig. 2 A). Deflections radially away from the center of the nucleus are displayed in yellow. In Fig. 2 B the radial deflection from 70 interphase cells were collected (4928 pillar deflections). The radial position for each pillar was normalized to the radius of each of the cell nuclei. The radial deflections, up to the radius of the nuclei (Fig. 2B inset, blue line), were symmetrically spread around Δ*x*/*R* = 0.005 ± 0.011 (mean ± standard deviation). Such spreading was predicted from the accuracy by which the center-of-mass of each pillar was determined. The distribution of radial deflections beyond the nuclei significantly widens, with Δ*x*/*R* = 0 ± 0.02 (Fig. 2 B inset, red line). Negative deflections seen below and around the nucleus, mainly reflect the inward-pointing traction forces that cells apply on their substrate. The cell shown in Fig. 2 pulled inward on the pillar array resulting in a maximum pillar deflection of 488 nm which is translated into a force of 6.7 nN.

**Fig. 2.**
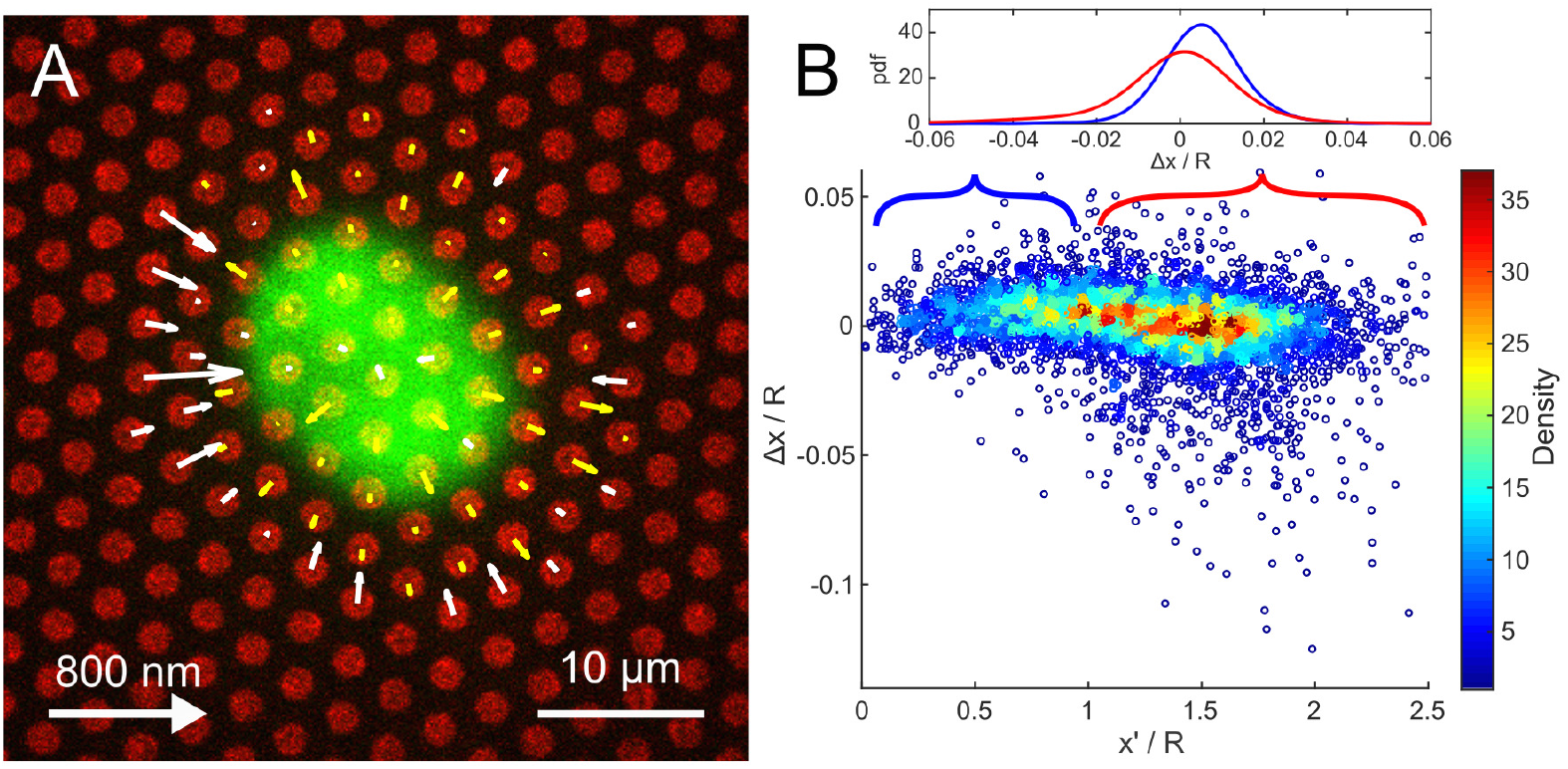
Nuclei of interphase cells induce astigmatism of measurable size, but do not influence the measurement of traction forces near the cell periphery. **A.** A HeLa cell during interphase on a 9.8 kPa micropillar array. The radial deflection of each pillar is shown by an arrow. We see the cell pulling pillars inward (white). Outward deflections are displayed in yellow. **B.** Relative displacement of dots underneath 70 interphase cells on soft micropillars. For *x′* < *R*, displacements are predominantly caused by astigmatism and we found Δ*x*/*R* = 0.005 ± 0.011 (inset, blue line). At *x′* > *R*, cells exerted traction forces on the pillars. The displacement was Δ*x*/*R* = 0 ± 0.02 (inset, red line). Values indicate mean ± SD.

### 3.4. Astigmatism induced aberrations change dynamically during mitosis

During cell division, cells undergo dramatic morphological and shape transitions. In metaphase, cells re-model their adhesion to the substrate [22], round up and take on a largely spherical shape [23]. We analyzed how the cell roundup during mitosis leads to apparent deflections governed in large by a change in astigmatism.

Figure 3 A-C shows snapshots of a 15-hour movie of the cell cycle (also see supplementary video). HeLa cells were transfected with a GFP-labeled histone-2B plasmid, and placed on a stiff (47.2 kPa) micropillar array. Radial pillar deflections beyond background deflections (see Methods) are shown by arrows. Inward deflections are displayed in white, outward deflections are shown in yellow. Analogous to Fig. 2, during interphase (Fig. 3 A) the cell and nucleus were spread over the micropillars. The cell exerted significant traction forces on some pillars, visible by the inward pulling of pillars at the cell perimeter (not visible). When the cell rounded up during metaphase (Fig. 3 B), large outward deflections close to the nucleus were observed. For quantitative analysis we calculated the mean outward radial deflection by averaging on all radial components of outward pointing deflections that were beyond background (Fig. 3 D). During cell division, the outward deflection per pillar increased from ~ 50 nm in interphase to ~ 120 nm in metaphase. Outward deflections were maximal between 470 and 520 min of the movie. Shortly after DNA separation at 520 min, outward deflections close to the nucleus were still significant. Subsequently, the two daughter cells again spread out on the substrate resulting in a gradual decrease in the net radial deflection to ~ 60 nm (Fig. 3 C-D).

**Fig. 3.**
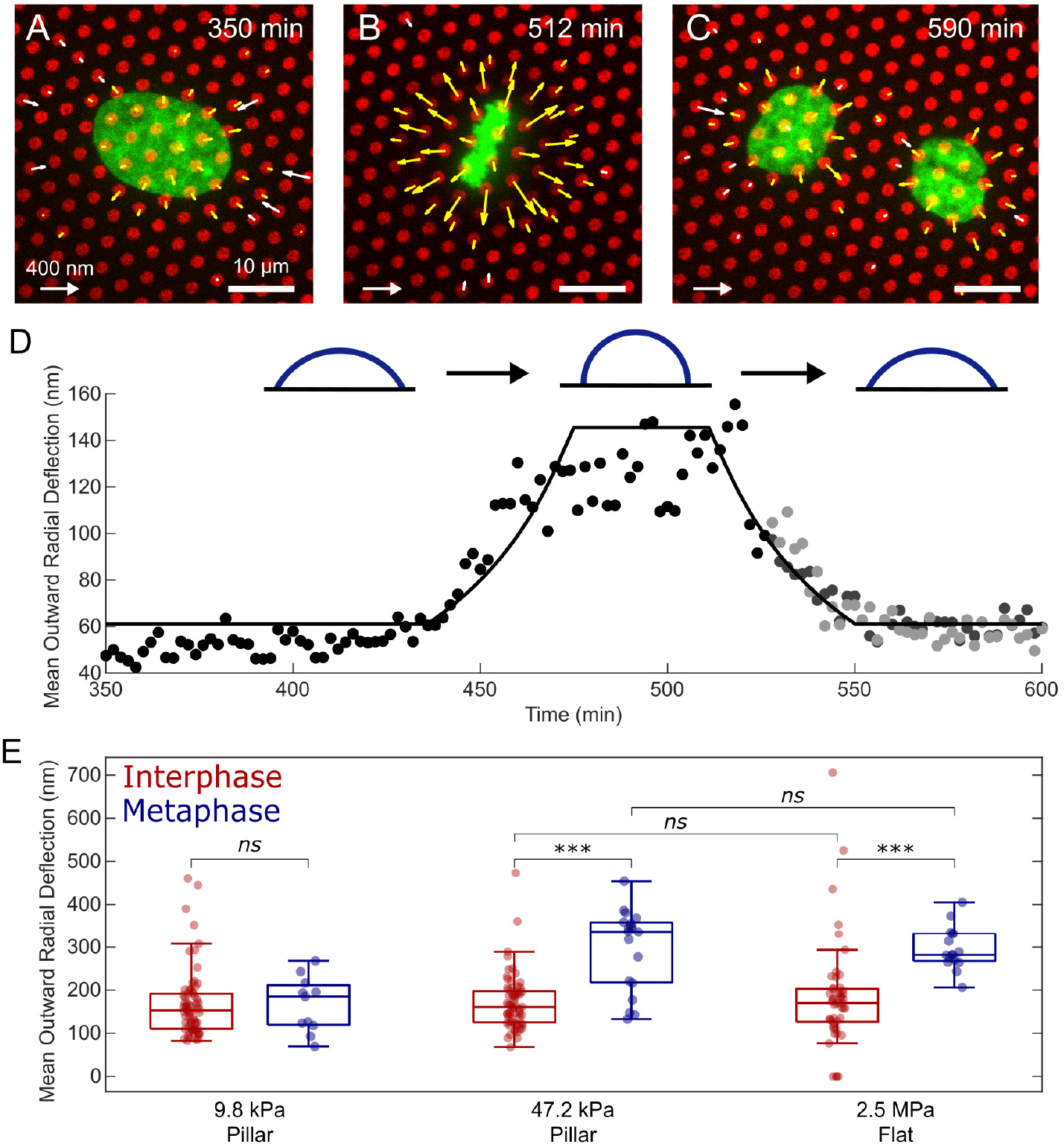
Apparent outward deflections significantly increase during metaphase. **A-C.** Snapshots of a HeLa cell during cell division on a 47.2 kPa micropillar array. Pillar tops are shown in red, the nucleus in green. The radial deflection of each pillar beyond the background is shown by an arrow. We see the cell pulling pillars inward (white). Outward deflections are displayed in yellow. The deflection scalebar in the lower left and fluorescent scalebar in the lower right are consistent for each image. **D.** Time series of the mean outward radial deflection of the cell in (A-C). The mean radial deflection had a peak-plateau during metaphase at around 475-515 min. After cell division (split into grays for each individual cell) the mean radial deflection gradually decreased. The solid line shows the expected mean deflection caused by a sphere-cap transitioning from flat to round and back based on Eq. (1). **E.** On the stiffer substrates, the mean outward radial deflection was higher during metaphase than during interphase. There is no statistical significant difference between stiff pillars and the flat substrate. From left to right 70, 11, 65, 19, 49, 16 cells were analyzed. *ns*: *p* > 0.05, ***: *p* < 0.001.

The increase in apparent radial deflection is described by the shape change of the nucleus from flat to round and back (inset Fig. 3 D). We modeled the shape change by a linear transition from flat to round and and back. Subsequently we applied the spherical cap-model (Eq. (1)) to calculate the predicted astigmatism. In our model we further assumed that the nuclear volume and its refractive index were constant. Based on Eq. (1), deflections were calculated and averaged over the area covered by the spherical-cap (solid black line in Fig. 3 D). For interphase cells (< 440 min) we set the height *H* = 8.0 μm, and radius *R* = 19.2 μm. Between 440 min and 475 min, the height increased to its maximum value in metaphase, with *H* = 9.4 μm, and the radius decreased to *R* = 8.6 μm. That shape change was followed by an increase of the mean apparent deflection from 61 to 146 nm. The cell’s rounded shape was conserved for 35 min, when mitosis finished. Subsequently, the cap-height decreased between 510 and 540 min, during which the mean apparent deflection reduced back to 61 nm. The change in apparent deflection, as observed in the experiments, was faithfully described by our simple astigmatism model coupled to the shape change during cell division. Hence, pillar displacements were not the result of a change in cellular traction but rather due to a change in astigmatism.

To further substantiate our findings we followed the behavior of cells on surfaces of various stiffness. The apparent radial deflection for 70 interphase and 11 metaphase cells on soft pillars of 9.8 kPa, 65 interphase and 19 metaphase cells on stiff pillars of 47 kPa, and 49 interphase and 16 metaphase cells on a continuous layer of PDMS of 2.5 MPa were analyzed. The results are summarized in Fig. 3 E. The mean outward pointing radial deflection per pillar was calculated for each cell. On 9.8 kPa pillars, no significant difference was found for cells in interphase and metaphase (*p* = 0.7), with deflection of 167 ± 79 nm and 167 ± 64 nm respectively. On stiff pillars of 47.2 kPa, we found a significant increase in apparent outward deflections for cells in metaphase as compared to interphase (*p* < 0.001). The mean deflection was 172 ± 65 nm and 297 ± 95 nm respectively. On the coated flat substrate, a similar trend was found. The mean deflection increased from 184 ± 124 nm in interphase to 297 ± 50 nm in metaphase (*p* < 0.001). More importantly, we found no significant difference between the results on stiff pillars with those on flat PDMS for cells in their respective stages of cell division (interphase *p* = 0.8, metaphase *p* = 0.5). These results indicate that all outward deflections observed were caused by astigmatism and not by cells (actively) pushing the pillars outwards. All p-values are summarized in supplementary table S1.

The negative radial deflections resulting from inward pointing traction forces that cells in interphase exerted on the substrate are shown in the supplementary Fig. S5. For the lowest stiffness substrate of 9.8 kPa the mean deflection was −218 ± 120 nm, which translates into a force of −3.0 ± 1.6 nN. Such forces are typical for adherent cells as earlier reported [24]. The deflections for high-stiffness pillars (47.2 kPa) and for the continuous substrate were significantly lower and indistinguishable from random deflections (*p* = 0.2) as of the positional accuracy for the detection of all pillars. This results in an upper limit of the mean traction force per deflected pillar of −8 ± 5 nN for HeLa cells.

## 4. Discussion

In our study we were concerned with the effect of cellular astigmatism in quantitative high-resolution imaging. We predicted that the presence of large curved objects, like the nucleus of the cell, will result in significant outward-pointing deflections when objects are imaged through the cell.

For quantification we showed that a simple model that approaches the shape of the nucleus to that of a spherical-cap was sufficient to describe the experimental results. A fit of the data to the model allowed us to obtain a value for the refractive index of the nucleus of ~ 1.369. Our value corroborates values reported earlier using other techniques [14].

The importance to take cellular astigmatism into account became apparent when we studied adherent cells on structured surfaces during mitosis. We found that apparent deflections increased up to 400 nm that misleadingly could be interpreted in cellular outward pushing forces during mitosis. However those apparent deflections quantitatively followed our predictions for astigmatism for cells that round-up during mitosis.

For practical use we defined a figure-of-merrit, *Z*, that compares the astigmatic displacements to the nominal resolution of a technique. Taking the refractive index of the nucleus that we report here, we showed that large objects in a cell like the nucleus will lead to barely visible effects in regular microscopy, *Z* ≈ 1. Yet for any high- or super-resolution technique astigmatism will be substantial, *Z* ≫ 1. In super-resolution techniques, for which resolutions of 30 nm are typically achieved, *Z* < 1 holds for objects smaller than 0.8 μm. Hence, the effect of astigmatic deformations for objects like vesicles [1] and bacteria [25] can be neglected.

It should be mentioned that astigmatism may be circumvented by carefully matching the refractive index of the embedding medium. This method has been successfully developed and implemented for cells [26]. Yet it is unclear how to implement such a strategy for multiple compartments, the cytosol and the nucleus, without breaking membranes allowing for life cell experimentation.

The figure-of-merit we defined here is a simple and robust criterion to judge whether astigmatism needs to be accounted for in a given experimental setting.

## Funding

This work is part of the research program “The Active Matter Physics of Collective Metastasis” with project number Science-XL 2019.022, which is financed by the Dutch Research Council (NWO).

## Acknowledgments

HH, CB, JE and TS designed the experiments. RRM and MV conducted the experiments. RRM performed data analysis. RRM and TS wrote the manuscript, all authors reviewed and edited the manuscript. JE and TS supervised the work.

## Disclosures

The authors declare no conflicts of interest.

## Notes

### Competing Interest Statement

The authors have declared no competing interest.

