## Supplemental Document for "Characterization of Cell-Induced Astigmatism in High-Resolution Imaging"

### Characterization of Cell-Induces Astigmatism in High-Resolution Imaging: Supplemental Document


#### 1. THEORETICAL FRAMEWORK

At the interface between two media with different refractive indices, light gets refracted. The curvature of the interface and the refractive index difference determine the strength of the lens. Because of the optical techniques used to image biological cells, cells qualify as lenses since they meet the requirements. First of all, nuclei of cells have a different refractive index than their surrounding [1]. Furthermore, during the cell cycle of mitotic animal cells, the shape of the cells changes considerably from flat to round [2]. The combination of these two make the cell a potential lens, especially during cell division. Here, we present a simple model to calculate astigmatism due to refraction of light at the interface between the nucleus and its surrounding.

##### A. Mitotic Cell as a Lens

As an approximation, we model the nucleus as a spherical-cap of radius $R$ and height $H \leq 2R$ above the surface of a substrate. Fig. S1 shows a cut of the sphere in 2D. The sphere is placed in a medium with refractive index $n_1$. The sphere itself has refractive index $n_2$. A light ray (red in Fig. S1) is refracted at the interface between the medium and the sphere. Due to refraction, a net shift $\Delta x$ is observed. To find the analytical expression of the shift, we start by relating the angle of refraction of the ray to the position on the sphere

$$\sin\theta_2 = \frac{x}{R}\,, \tag{S1}$$

where $\theta_2$ is the incident angle at position $x$.

Using Snell's law we can find an expression for the incident angle $\theta_2$ in terms of geometrical constants

$$\sin\theta_1 = \frac{n_2}{n_1}\sin\theta_2 = \frac{n_2}{n_1}\frac{x}{R}\,. \tag{S2}$$

The difference $\Delta x$ between the observed position $x'$ and the real position $x$ is

$$\frac{\Delta x}{R} = \frac{x' - x}{R} = \frac{h}{R}\tan(\theta_1 - \theta_2)\,, \tag{S3}$$

where $h$ is the height of the sphere at position $x$.

Defining $L = h + R - H$, and realizing that $R$ and $x$ are normal such that $L^2 = R^2 - x^2$ an expression for $h/R$ is found

$$\left.\begin{array}{l} \frac{L}{R} = \frac{1}{R}\sqrt{R^2 - x^2} = \sqrt{1 - \frac{x^2}{R^2}} \\ L = h + R - H \end{array}\right\} \frac{h}{R} = \sqrt{1 - \frac{x^2}{R^2}} - \left(1 - \frac{H}{R}\right)\,. \tag{S4}$$

Then, combining Eq. (S3) and Eq. (S4) results in

$$\frac{\Delta x}{R} = \left[\sqrt{1 - \left(\frac{x}{R}\right)^2} - \left(1 - \frac{H}{R}\right)\right]\tan\left(\theta_1 - \theta_2\right)\,. \tag{S5}$$

Further, combining Eq. (S1), Eq. (S2) and Eq. (S5) results in the final expression for the apparent deflection $\Delta x$

$$\Delta x = R \left[ \sqrt{1 - \left(\frac{x}{R}\right)^2} - \left(1 - \frac{H}{R}\right) \right] \tan \left( \arcsin \frac{n_2}{n_1} \frac{x}{R} - \arcsin \frac{x}{R} \right) , \tag{S6}$$

which, as a function of $x$, in the end only depends on the radius of the sphere $R$, the height $H$ that is above the surface, and the refractive indices of the medium $n_1$ and the sphere $n_2$. For typical values ($R = 10$ m, $H = 7$ m, $n_1 = 1.33$ and $n_2 = 1.38$) aberrations close to the edge of the sphere can be up to 300 - 700 nm.

Approximating for distances close to the center, i.e. $x << R$, we find a linear dependency of the lateral displacement with the distance from the center of the sphere

$$\frac{\Delta x}{R} \approx \frac{H}{R} \left( \frac{n_2}{n_1} - 1 \right) \cdot \frac{x}{R} . \tag{S7}$$

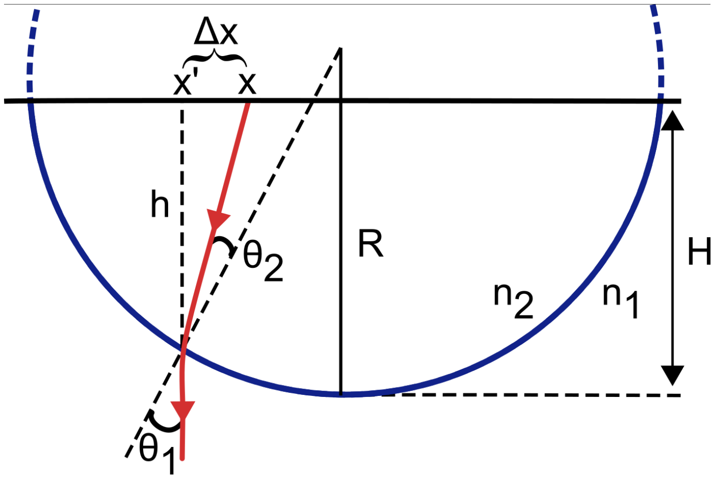


**Fig. S1. Refraction at a spherical surface results in a shift $\Delta x$ between the real pillar position $x$ and the observed pillar position $x'$.** A light ray (red) is shown for the case that $n_2 > n_1$.

#### 2. SUPPLEMENTAL FIGURES

**Table S1.** P-values resulting from a Dunn's test pairwise comparison after a Kruskal-Wallis test of the distributions of mean outward radial deflections (Fig. 3). Distributions are from each 9.8 kPa micropillars, 47.2 kPa micropillars and a 2.5 MPa flat PDMS layer of cells in interphase (I) and metaphase (M).

| | | 9.8 kPa | | 47.2 kPa | | 2.5 MPa | |
|---|---|---|---|---|---|---|---|
| | | I | M | I | M | I | M |
| 9.8 kPa | I | 1 | 0.7 | 0.4 | $< 0.001$ | 0.3 | $< 0.001$ |
| | M | 0.7 | 1 | 0.9 | 0.002 | 0.8 | $< 0.001$ |
| 47.2 kPa | I | 0.4 | 0.9 | 1 | $< 0.001$ | 0.8 | $< 0.001$ |
| | M | $< 0.001$ | 0.002 | $< 0.001$ | 1 | $< 0.001$ | 0.5 |
| 2.5 MPa | I | 0.3 | 0.8 | 0.8 | $< 0.001$ | 1 | $< 0.001$ |
| | M | $< 0.001$ | $< 0.001$ | $< 0.001$ | 0.5 | $< 0.001$ | 1 |

**Table S2.** P-values resulting from a Dunn's test pairwise comparison after a Kruskal-Wallis test of the distributions of mean inward radial deflections (Fig. S5). Distributions are from each 9.8 kPa micropillars, 47.2 kPa micropillars and a 2.5 MPa flat PDMS layer of cells in interphase (I) and metaphase (M).

| | | 9.8 kPa | | 47.2 kPa | | 2.5 MPa | |
|---|---|---|---|---|---|---|---|
| | | I | M | I | M | I | M |
| 9.8 kPa | I | 1 | $< 0.001$ | $< 0.001$ | $< 0.001$ | 0.006 | $< 0.001$ |
| | M | $< 0.001$ | 1 | 0.001 | 0.7 | $< 0.001$ | 0.04 |
| 47.2 kPa | I | $< 0.001$ | 0.001 | 1 | $< 0.001$ | 0.2 | 0.4 |
| | M | $< 0.001$ | 0.7 | $< 0.001$ | 1 | $< 0.001$ | 0.05 |
| 2.5 MPa | I | 0.006 | $< 0.001$ | 0.2 | $< 0.001$ | 1 | 0.08 |
| | M | $< 0.001$ | 0.04 | 0.4 | 0.05 | 0.09 | 1 |

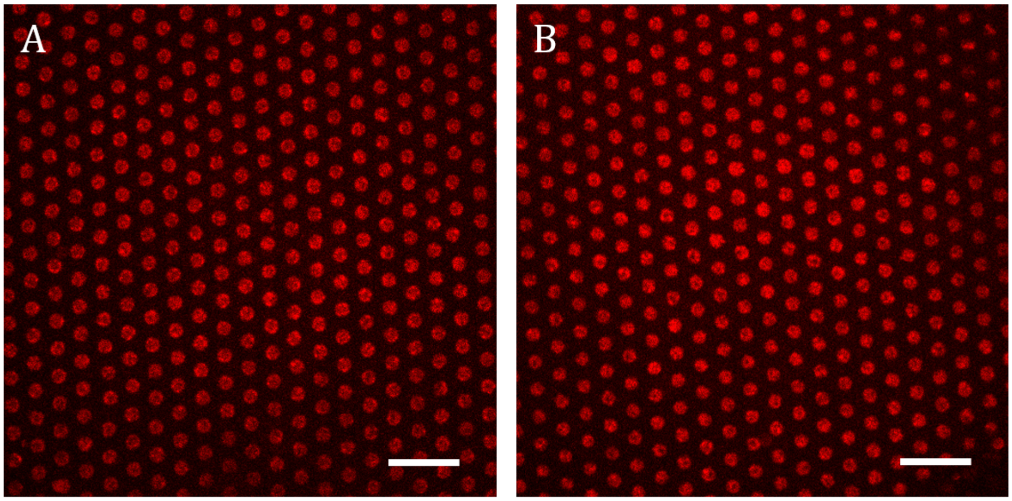


**Fig. S2. Result of micro-contact printing of fibronectin on a micropillar array (A) and through a two-step stamping method on a flat layer of PDMS (B).** The scalebar in the lower right is 10 m.

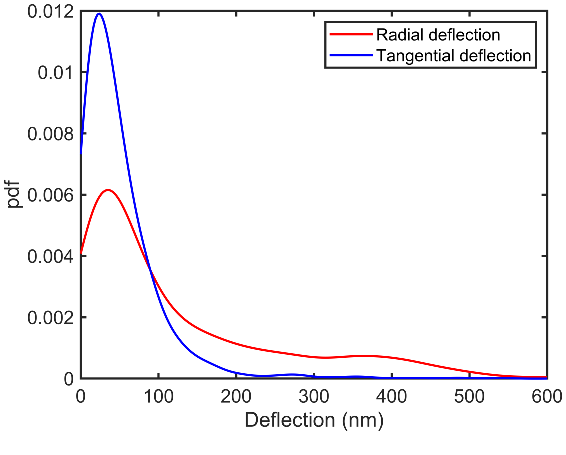


**Fig. S3. Radial deflections were much larger compared to tangential deflections.** Close to the nucleus, astigmatism dominates the random deflections. The apparent deflection caused by astigmatism are pointing radially away from the center of the nucleus. The tangential component of the pillar deflections were mainly caused by inaccuracies in the pillar-centroid determination or inhomogeneities of the array.

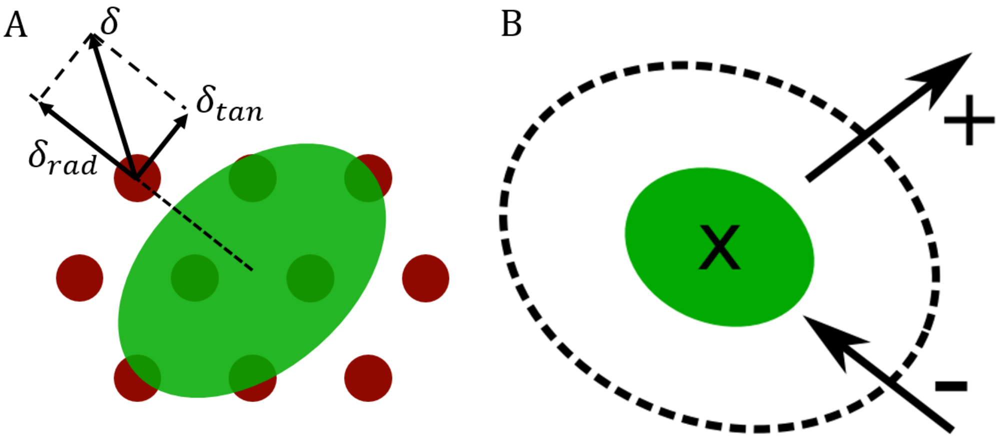


**Fig. S4. Definition of radially in- and outward deflected pillars. A.** Deflections $\delta$ of pillars (red) were separated into radial $\delta_{rad}$ and tangential $\delta_{tan}$ components with respect to the center of the nucleus (green). **B.** Deflections radially outward from the center of the nucleus were defined as positive, inward deflections were defined as negative. (Dotted line marks cellular outline, x marks relative center for radial deflections)

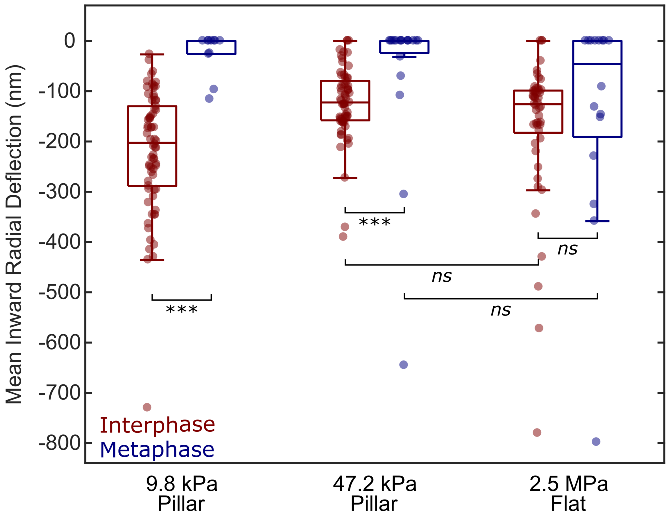


**Fig. S5. Inward deflections on stiff pillars were indistinguishable from deflections on the flat substrate.** In general, no inward deflected pillars were found during metaphase, suggesting cells fully detach from the substrate during this stage of cell division. On the softest, 9.8 kPa, micropillars, interphase cells were able apply sufficient force to bend pillars beyond the background. The mean traction force applied was $-3.0 \pm 1.6$ nN (mean $\pm$ SD) per deflected pillar. Although inward deflections were observed for interphase cells on stiff 47.2 kPa pillars (corresponding to $-8 \pm 5$ nN), they were not distinguishable from from random deflections ($p = 0.2$).
